# *In vivo* inducible reverse genetics in patients’ tumors to identify individual therapeutic targets

**DOI:** 10.1101/2020.05.02.073577

**Authors:** Michela Carlet, Kerstin Völse, Jenny Vergalli, Martin Becker, Tobias Herold, Anja Arner, Wen-Hsin Liu, Veronika Dill, Boris Fehse, Claudia D. Baldus, Lorenz Bastian, Lennart Lenk, Denis M. Schewe, Johannes W. Bagnoli, Philipp J. Jost, Cornelius Miehting, Kristoffer Riecken, Marc Schmidt-Supprian, Vera Binder, Irmela Jeremias

## Abstract

High-throughput sequencing describes multiple alterations in each tumor (1), but their functional relevance is often unclear. Clinic-close, individualized molecular model systems are required for functional validation and to identify therapeutic targets of high significance for each patient (2). Here, we established a Cre-ER^*T*2^-loxP based inducible RNAi-mediated gene silencing system in patient-derived xenograft (PDX) models of acute leukemias *in vivo*. Mimicking anti-cancer therapy in patients, gene inhibition was initiated in mice harboring orthotopic tumors. Fluorochrome guided, competitive *in vivo* trials identified a major tumor-maintaining potency of the MLL-AF4 fusion protein and validated MCL1 as vulnerability in some, but not all patients’ tumors. We could prove DUX4 to play an essential role in patients’ leukemias carrying the recently described DUX4-IGH translocation. By individualizing functional genomics in established tumors *in vivo*, our technique decisively complements the value chain of precision oncology. Being broadly applicable to tumors of all kinds, it will considerably reinforce personalizing anti-cancer treatment in the future.

## Main Text

Translating comprehensive cancer sequencing results into targeted therapies has been limited by shortcomings of model systems and techniques for preclinical target validation. The methodological gap contributes to the fact that only below 10% of drugs, successful in preclinical studies, pass early clinical evaluation and receive approval (3, 4). Functional genomic tools including RNA interference (RNAi) proved of utmost importance to annotate the numerous alterations detected by multi-omics profiling and significantly deepened our understanding of the merit of individual genes as drug targets (5, 6). As limitation, functional studies have largely been restricted to cancer cell lines, which often fall short in predicting the role of alterations in individual human tumors (7). To approximate the situation of the patient, the predictive power of primary tumor cell cultures (8) and organoids (9) is currently under intense investigation (10). For mirroring the clinical situation even closer, patient derived xenograft (PDX) mouse models have been demonstrated to faithfully recapitulate the complexity of tumors in humans. PDX models are available for the vast majority of human cancers, and their preclinical value for biomarker identification and drug testing is well established (11). It is increasingly recognized that the drug development process might profit from studying PDX models with molecular techniques, routinely used in cell line models and genetically engineered mouse models (GEMM) (12, 13). Still, RNAi techniques were only rarely applied for *in vivo* mechanistic studies in PDX, while inducible systems are entirely missing. As an advantage over constitutive systems, inducible gene silencing prevents over-estimating *in vivo* gene function by avoiding influences from, e.g., transplantation and engraftment, and allows mimicking the treatment situation in patients with established tumors. Here, we report the first inducible RNAi in PDX models *in vivo*, using acute leukemia (AL) as prototype disease where ex-vivo investigation on primary cells is challenging, but orthotopic PDX models are promising (14, 15).

Primary tumor cells from 7 patients with AL [3 pediatric acute lymphoblastic leukemia (ALL), 1 adult ALL, 3 adult acute myeloid leukemia (AML)] were transplanted into NOD scid gamma (NSG) mice (Figure 1a, clinical patient data in *Table S1*). Resulting PDX cells were lentivirally transduced first with a construct encoding a Tamoxifen (TAM)-inducible variant of Cre-recombinase, Cre-ER^*T*2^, together with a red fluorochrome for enriching transgene expressing cells and Gaussia luciferase (Luc) for *in vivo* imaging (16) (Figure 1a). A single vector integration per cell genome was achieved, ensuring homogenous expression levels of Cre-ER^*T2*^ (*Figure S1a*), minimal toxicity and neglectable leakiness in all samples, thus overcoming one of the challenges of TRE-based inducible expression systems (13).

**Fig. 1.**
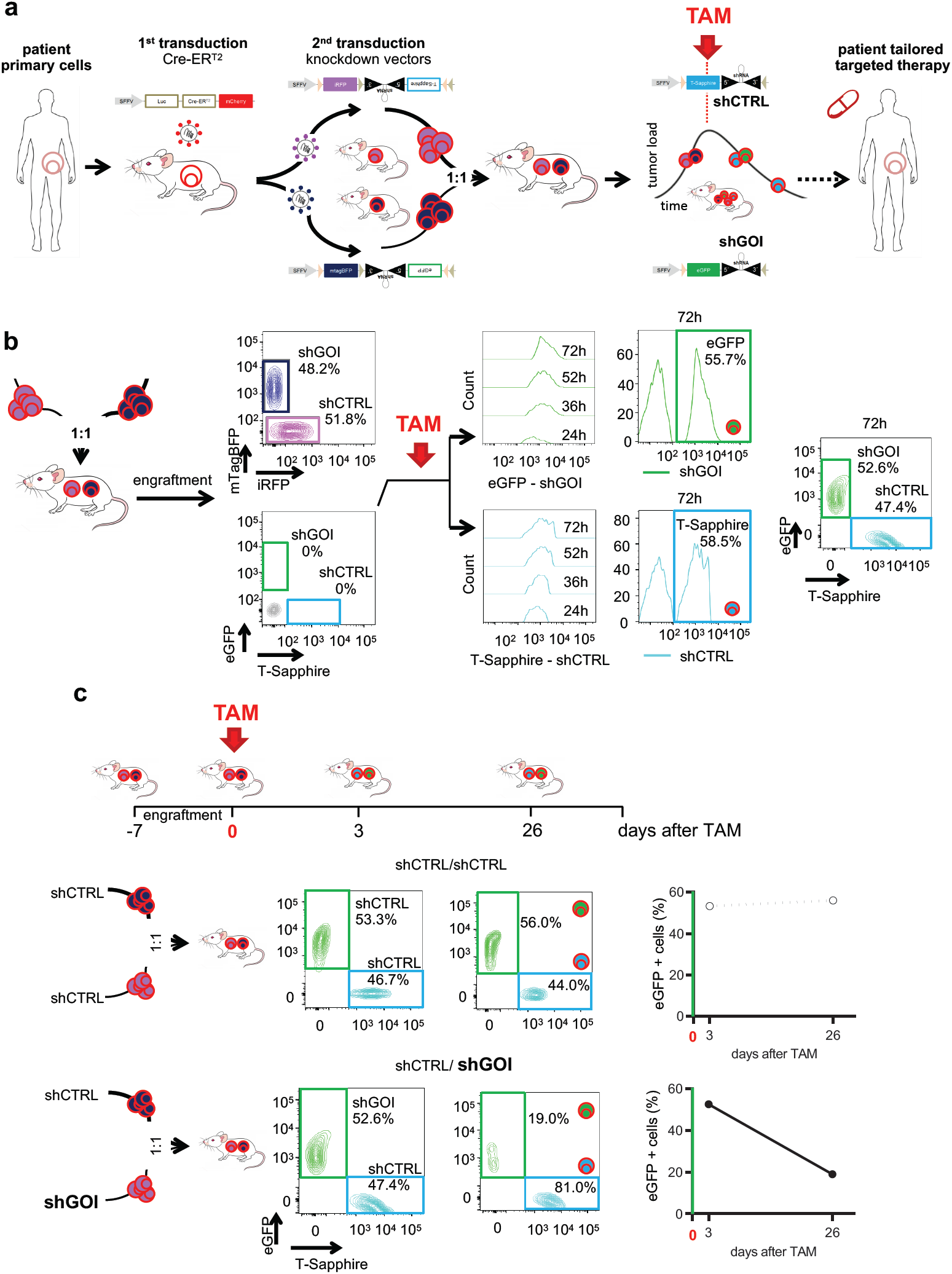
Establishing an inducible knockdown system in PDX acute leukemia cells *in vivo*. **a** <u>Overview of the experimental setup:Primary</u> acute leukemia (AL) cells were amplified in NSG mice and resulting PDX cells lentivirally transduced twice in a row; first to constitutively express Cre-ER^*T*2^ together with mCherry and a luciferase (Luc); second to express inducible knockdown vectors containing (i) a constitutively expressed fluorochrome marker (either iRFP or mTagBFP) and (ii), placed in antisense orientation, a miR30-based knockdown (KD) cassette coupled to a second inducible fluorochrome (either T-sapphire or eGFP). After amplification in mice, purified transgenic PDX cells were mixed 1:1 and transplanted into next recipient mice for competitive *in vivo* experiments. In mice with established leukemias, TAM was administered to induce Cre-ER^*T*2^ -mediated recombination. Recombination inverted the KD cassette and induced i) expression of the shRNA; ii) deletion of the constitutive fluorochrome (either iRFP or mTagBFP) and iii) expression of the inducible fluorochrome (either T-Sapphire or eGFP; see *Figure S1b* for detailed description). As result, T-Sapphire positivity indicated cells expressing the shRNA targeting a control (shCTRL), while eGFP positivity indicated cells expressing the shRNA targeting the gene-of-interest (shGOI). If the GOI harbors an essential function, the eGFP-positive population gets lost over time *in vivo*, indicating that the patient might profit from drugs targeting the GOI. **b** <u>Switch in fluorochrome expression upon Cre-ER*T*^2^-recombination:</u> Double transgenic PDX AML-388 cells expressing Cre-ER^*T*2^ together with either iRFP/shCTRL or mTagBFP/shGOI (shMCL1) were mixed 1:1 and 300.000 cells were injected into the tail vein of NSG mice (n=14). 7 days after injection, 2 mice were taken down and PDX cells analyzed by flow cytometry for all 4 fluorochromes. In the remaining mice, 50 mg/kg TAM was administered by oral gavage to induce Cre-ER^*T*2^ -mediated recombination. Resulting increase in T-Sapphire or eGFP expression, indicating expression of shCTRL and shGOI, respectively, was measured in PDX cells isolated from mice at the indicated time points (24, 36, 52 and 72h after TAM; n=3 per time point). Representative histograms and plots are shown. **c** <u>Typical results for a GOI with essential function:</u> Upper scheme depicts the experimental procedure: For pairwise competitive assays, mice were injected with either of two mixtures: a control mixture of iRFP/shCTRL and mTagBFP/shCTRL (short shCTRL/shCTRL) or the experimental mixture iRFP/shCTRL and mTagBFP/shGOI (short shCTRL/shGOI). TAM was administered 7 days after injection (day 0). Mice were sacrificed 3 and 26 days after TAM and PDX cells analyzed for expression of the inducible fluorochromes T-Sapphire and eGFP. Density plots show representative results for both mixtures on day 3 (left) and day 26 (right). Right panels show a quantification as percentage of [eGFP/shGOI positive cells divided by (the sum of T-Sapphire/shCTRL positive plus eGFP/shGOI positive cells)]; the shCTRL/shCTRL mixture is analyzed and depicted, respectively.

In a second step, PDX cells were transduced with the small hairpin (sh) RNA expression vectors (Figure 1a and *S1b*). The miR30-based knockdown cassette was directly coupled to a fluorochrome and both were cloned in antisense orientation, flanked by two pairs of loxP sites. In the absence of TAM, neither the inducible fluorochrome nor the shRNA were expressed. TAM administration induced a two-step Cre-ER^*T*2^-mediated recombination process which flipped the fluorochrome-shRNA insert into sense orientation, initiating its expression (*Figure S1b*) (17, 18). A set of 4 recombinant fluorochromes was used to monitor shRNA transduction and recombination and to enable competitive *in vivo* assays. Transduction efficiency was tracked by iRFP in a control vector carrying a shRNA targeting Renilla luciferase (shCTRL) or mTagBFP in the vector targeting the gene of interest (GOI). Upon TAM administration, CreER^*T*2^-mediated recombination deleted the constitutively expressed fluorochromes iRFP and mTagBFP and induced expression of the second set of fluorochromes (19) (Figure 1a and *S1b*). T-Sapphire and eGFP were chosen as inducible fluorochromes due to their high similarities in sequence and expression kinetics (20) and replaced iRFP and mTagBFP expression upon TAM treatment. The two knockdown vectors enabled pairwise competitive *in vivo* experiments in the same animal to increase reliability and sensitivity, while saving resources.

Mice were transplanted with a 1:1 mixture of PDX cells expressing either of the two RNAi vectors, shCTRL or shGOI (Figure 1a). As quality control, expression of constitutive markers revealed equal engraftment of both populations (Figure 1b). To induce gene silencing, TAM was given to mice with pre-established leukemias, mimicking treatment of patients with pre-existing tumors. Upon induction, the functional consequences of control and GOI knockdown were monitored by quantifying each population according to their fluorochromes, using flow cytometry (Figure 1b-c). Systemic TAM administration induced expression of the inducible fluorochromes T-Sapphire or eGFP, in similar amounts for both constructs, starting as early as 24h hours after TAM (Figure 1b). TAM was dosed to obtain substantial Cre-ER^*T*2^-induced recombination in the absence of toxicity. Recombination efficiency was found to be independent of tumor load, indicating that our model allows studying gene function at all disease stages (*Figure S1c*). Several quality controls were performed to exclude unspecific toxicities; the distribution of both populations remained stable over time after TAM treatment, if both populations expressed shCTRL (shCTRL/shCTRL mixture in Figure 1c, upper lane; representative for identical results in all samples). Similarly, the distribution of the shCTRL/shGOI mixture remained unchanged, if mice received the carrier solution alone (*Figure S1d-e*). In contrast and upon treatment with TAM, the population expressing a shRNA targeting an exemplary essential GOI decreased over time and was overgrown by control cells (Figure 1c, lower lane and (*Figure S1f*). Loss of eGFP positive shGOI cells started early, before the expression of the inducible fluorochrome T-Sapphire in the shCTRL group had reached its maximum expression level (*Figure S1f*). Loss of cells with GOI knockdown *in vivo* proved a functional importance of the GOI on the molecular level, mimicking elimination of tumor cells in patients upon treatment with a drug inhibiting the GOI.

To test the applicability of our approach, we started studying a bona fide positive control with high likelihood of harboring an essential function in established PDX tumors *in vivo*. The translocation t(4;11) leading to the expression of the MLL-AF4 fusion (KMT2A-AFF1) is present in 80% of infant B-precursor ALL patients, and is associated with poor prognosis (21). Several studies elucidated its role in ALL cell lines and mouse models (22), but up to date no molecular investigations on its function have been carried out in patient cells or established tumors growing *in vivo*. We designed a shRNA targeting a mRNA breakpoint shared by several patients, which was able to reduce expression of the fusion transcript (Figure 2a); as the shRNA sequence targeted neither of the individual wildtype genes, MLL or AF4, it might induce minor adverse effects on normal tissue when applied *in vivo*, e.g., by systemic gene therapeutic approaches (Figure 2a). Inducible knockdown of MLL-AF4 significantly reduced ALL cells in the t(4;11)-positive PDX model tested, proving a tumor maintaining role of MLL-AF4 in established patient tumors *in vivo*, while no effect was observed in the competitive experiment where a mixture of shCTRL/shCTRL cells was injected (Figure 2b). Variations between the different animals were neglectable reflecting the high reliability of our approach (Figure 2b). Reduced tumor growth of the shMLL-AF4 mixture was visible using *in vivo* imaging, even though 50% of injected tumor cells were shCTRL cells (Figure 2c). Non-MLL-AF4 rearranged ALL remained unaffected (*Figure S2a*), demonstrating the specificity of our approach. These results prove the selectivity and operability of our technique and showed for the first time that MLL-AF4 harbors an essential function in established patient-derived leukemias growing *in vivo*. We provide strong molecular evidence that the translocation transcript represents an attractive therapeutic target for future therapies.

**Fig. 2.**
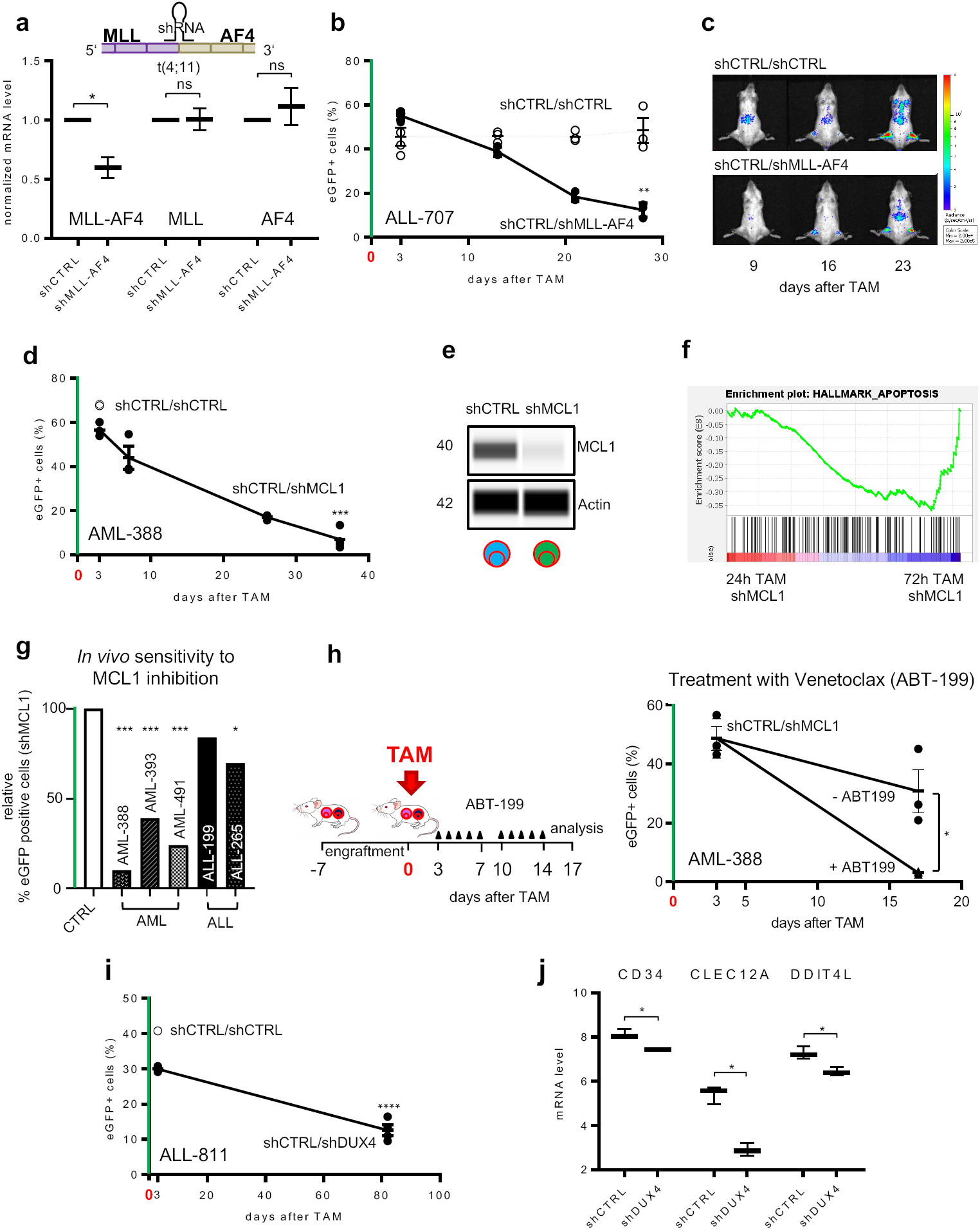
Identifying functionally relevant therapeutic targets in individual tumors *in vivo*. **a – c** MLL-AF4 plays an essential role *in vivo* in rearranged ALL.**a** A shRNA targeting the MLL-AF4 fusion mRNA was designed according to the patient-specific breakpoint of PDX ALL-707 (*Table S2*). mRNA expression of MLL-AF4, MLL and AF4 was analyzed by qPCR in CTRL and MLL-AF4 knockdown ALL-707. Mean±SEM of cells isolated from 4 mice 28 days after TAM are shown.*p<0.05, Welch’s t-test; ns not significant. **b** Competitive experiments were performed and analyzed as in Figure1c, using PDX ALL-707 cells and the shGOI targeting MLL-AF4. TAM was applied on two consecutive days (100 mg/kg, day -1+0). Mice were sacrificed 3 (n=4), 13 (n=3), 21 (n=3) and 28 (n=3) days after TAM; each dot represents one mouse; mean±SEM; *p<0.01, unpaired t-test. To determine significance of depletion of shGOI-expressing cells, the percentage of eGFP/shGOI cells at the experimental endpoint is compared to the percentage of eGFP/shCTRL cells at 72h post TAM (n=3), as this time point is used to define the sample-specific recombination efficiency. **c** Representative *in vivo* bioluminescence images of mice bearing a shCTRL/shCTRL or shCTRL/shMLL-AF4 mixture from the experiment described in Figure2b, at the indicated time points after TAM administration. **d – h** <u>MCL1 plays an essential role for some, but not all acute leukemia samples</u> **d** AML-388 PDX growing in mice rely on MCL1. Competitive experiments were performed and analyzed as described in Figure 2b, using PDX AML-388 cells and the shGOI targeting MCL1. TAM (50 mg/kg) was applied once (day 0); mice were sacrificed 3 (n=3), 7 (n=3), 26 (n=3) and 36 (n=4) days after TAM. Mean±SEM of the proportion of eGFP-positive cells isolated from shCTRL/shCTRL or shCTRL/shMCL1 bearing mice out of all recombined cells is displayed; each dot represents one mouse. ***p<0.001 by unpaired t-test. **e** Determination of MCL1 protein level. PDX AML-388 cells from the experiment in Figure 2d isolated 7 days after TAM and sorted for T-Sapphire/shCTRL and eGFP/shMCL1 were analyzed by immunoassay (Simple Western); *β*-Actin served as loading control; one representative example out of 3 replicates is shown; quantification in Figure S2b. **f** Gene set enrichment analysis (GSEA) of transcriptome data from cells of experiment in *Figure S2e*, isolated 24 and 72 hours after TAM and sorted for eGFP/shMCL1 (n=3 per time point). NES= -1.52, P=0.0. **g** Summary of results obtained in 5 PDX AL models with n=6 mice per group per PDX sample; experiments were performed and analyzed as described in Figure2d; shown is ratio of eGFP/shMCL1 cells normalized to eGFP/shCTRL at day 3 post-TAM. ***p<0.001,**p<0.01 by unpaired t-test. **h** The combinatorial effect of MCL1 KD plus ABT-199 (Venetoclax) was studied by injecting mice with a 1:1 mixture of shCTRL/shMCL1 AML-388 cells (300.000 cells/mouse) and treating them with 50 mg/kg TAM 7 days after injection (day 0). 72h after TAM administration, control mice were sacrificed (n=3) and the remaining mice injected i.p. either with 100 mg/kg ABT-199 (n=3) or PBS (n=3) for 5 consecutive days per week, in 2 cycles. At the end of the experiment (17 days after TAM), mice were sacrificed and analyzed as in Figure 1c. Mean±SEM is shown; *p<0.05 by unpaired t-test. **i**,**j** <u>DUX4 plays an essential role for DUX4-IGH rearranged ALL</u> **i** Competitive experiments were performed and analyzed as in Figure 2b, using ALL-811 and the shRNA targeting DUX4. 10 days after injection, TAM (50 mg/kg) was applied once (day 0). Mice were sacrificed 3 (n=2) and 82 (n=4 shDUX4) days after TAM. Shown is mean±SEM. ****p<0.0001 by unpaired t-test. **j** From the experiment in Figure 2i, transcriptome analysis was performed from eGFP/shCTRL and eGFP/shDUX4 cells 82 days after TAM (n=3 for each condition). Median and 95% CI of the three most differentially expressed genes are depicted; *p<0.05 corrected by multiple testing.

We next decided studying a gene where up to date and based on descriptive sequencing data, no predictive biomarker for response to target inhibition exists. The anti-apoptotic gene MCL1 is dysregulated in numerous tumor entities (23) and MCL1 inhibitors are currently investigated in clinical trials yielding mixed results (24) (NCT03218683). Predicting treatment response for selecting patients who will profit from MCL1 directed therapy remains a major challenge and functional *in vivo* assays might provide helpful insights (25). We studied PDX models from three different patients with acute myeloid leukemia (AML-388, 393, 491) and two patients with ALL (ALL-199, 265). In all AML PDX models, we found a clear decrease of cells with MCL1 knockdown compared to control cells *in vivo*, accompanied by efficient KD on protein level (Figure 2d-e, *Figure S2b and S2c-d*), validating MCL1 as important vulnerability. Silencing MCL1 induced rapid cell death, most prominent in the first 72 hours after administering TAM (*Figures S2c-e*). Gene set enrichment analysis from RNA sequencing data indicated that MCL1 knockdown was associated with activation of the apoptosis pathway, verified using Annexin-V staining (Figures 2f and *Figure S2f*). In addition, re-transplantation experiments into zebrafish (danio rerio) confirmed significant and rapid depletion of PDX cells upon MCL1 knockdown in an additional, independent *in vivo* model (*Figures S2g-i*).

In contrast to AML, inducible silencing of MCL1 in ALL samples showed minor to no effects, proving sub-entity specificity and patient-individual sensitivities (Figure 2g and *Figure S2j-k*). Functional relevance could not be predicted by expression levels of anti-apoptotic BCL-2 family members, highlighting the need for functional assays (*Figure S2l*). As MCL1 has been shown to confer resistance to several anticancer drugs (26), we next examined whether inhibition of MCL1 strengthens the response of AML PDX models towards drug treatment. We found that treatment with the BCL-2 inhibitor ABT-199 (Venetoclax) (Figure 2h) or the conventional chemotherapeutic drug Cytarabine (*Figure S2m*) further decreased the MCL1 knockdown population, indicating that sensitivity towards ABT-199 or Cytarabine might be increased by MCL1 directed treatment in patients. Thus, using MCL1 as exemplary target, we provide evidence that our approach enables distinguishing between subgroups of tumors in order to select patients, which might profit from therapies targeting a certain GOI, and to evaluate treatment combinations.

In a last step, we examined a less well studied tumor alteration with less defined function and selected the recently discovered rearrangement t(4;14) which occurs in 7% of ALL patients and results in the DUX4-IGH gene fusion (27). As cells with t(4;14) display high levels of otherwise absent DUX4, we asked whether DUX4 represents an important alteration in this subgroup of ALL. Our technique clearly revealed an essential function for DUX4 in t(4;14) rearranged ALL (Figure 2i). Expression of the DUX4-IGH translocation was reported to be associated with a defined gene expression signature (28, 29); we observed reversal of this signature upon silencing DUX4 in PDX models *in vivo* (Figure 2j and *Figure S2n*). Our technique could thus identify DUX4 as attractive therapeutic target to treat the recently detected subgroup of DUX4-IGH rearranged ALL.

In summary, our method evaluates the functional relevance of tumor alterations (i) in the background of individual patient tumors and their specific characteristics; (ii) in the complex *in vivo* environment of living beings and; (iii) in the situation of a pre-existing tumor, avoiding influences irrelevant for patients. Our molecular approach closely mimics the clinical situation and complements an important step in the evaluation chain of precision oncology. The molecular technique allows target validation independently from confounders such as pharmacodynamics and pharmacokinetics, toxicity and lack of specificity, inherent to drugs and compounds (30). The approach is devoid of overestimation by model-inherent processes like transplantation, homing and engraftment. Our knockdown approach might complement CRISPR/Cas9-mediated knockout approaches (13), while putatively more coherently mimicking the partial, but incomplete target inhibition induced by drugs or compounds. In addition to alterations detected by sequencing, our technique allows functional evaluation in an agnostic approach, e.g., in cell death pathways and studying “un-druggable” targets, including non-coding RNAs (31). While we studied acute leukemias as model diseases, the CRE-loxP-system was successfully used in numerous different tumor entities and our technique can easily be transferred to other cancers; our approach is thus relevant for tumor diseases of all kinds, far beyond leukemias. We envision a major potential of our method on a proof-of-concept level, where deeper knowledge on tumor dependencies will improve drug design and the ability to interpret patient sequencing data. It might also serve as a highly clinic-related, functional biomarker to improve clinical decision making to individualize treatment. Due to its major potential to tailor drug development, improve patient care and increase the success rate of clinical trials, our technique will foster personalized oncology in the future.

## Materials and Methods

### Ethical Statements

Written informed consent was obtained from all patients and from parents/carers in the cases where patients were minors. The study was performed in accordance with the ethical standards of the responsible committee on human experimentation (written approval by Ethikkommission des Klinikums der Ludwig-Maximilians-Universität München, April 15/2008, number 068-08; September 24/2010, number 222-10; January 18/2019, number 222-10) and with the Helsinki Declaration of 1975, as revised in 2000. All animal trials were performed in accordance with the current ethical standards of the official committee on animal experimentation (written approval by Regierung von Oberbayern, January 15/2016, Az. ROB-55.2Vet-2532.Vet02-16-7; Az. ROB-55.2Vet-2532.Vet02-15-193; ROB-55.2Vet-2532.Vet03-16-56).

### Cloning

For constitutive expression of the Cre-ER^*T*2^ recombinase, the coding sequence of the enzyme was PCR amplified from the CreERT2FrtNeoFrt cassette (gift from MSS) using a 5’ primer carrying NsiI and a 3’ primer carrying P2A-NsiI and ligated into the NsiI digested pCDH-SFFV-GLuc-T2A-mCherry vector downstream of the T2A peptide (*Figure S2a*) (pCDH-vector, System Bioscience). For inducible knockdown of target genes, the lentiviral FLIP vector system (17, 18) was optimized to link shRNA expression to fluorochrome expression. We used the lentiviral pCDH backbone, digested the vector with SpeI and SalI and introduced the following elements as a pre-synthetized stretch of DNA (GenScript®, Piscataway, NJ, USA): SpeI - SFFV - lox2272 - mtagBFP (iRFP720) - lox5171 - mir30 cassette-eGFP (T-Sapphire) -lox2272 - lox5171 – SalI (*Figure S2b*). The shRNA sequences targeting the different genes (MCL1, DUX4; see *Table S2*) were designed using the SplashRNA algorithm (32), with the exception of MLL-AF4 where sequences were designed to directly cover the patient-specific translocation breakpoint (*Table S2*). As control, a shRNA targeting the Renilla luciferase was used in all experiments (shCTRL). The shRNA-sequences were introduced into the miR30 cassette of the KD vector as part of pre-synthetized and annealed, complementary single strand DNA oligos (110 bps, see *Table S2*; Integrated DNA Technologies, USA), having XhoI and EcoRI as 5’ and 3’ restriction sites, respectively. For knockdown of MLL-AF4, the miR-E KD cassette was used (33) and concatemerized to enhance the knockdown efficiency (34).

### Generating transgenic PDX models

Establishing AML and ALL PDX models in NOD scid gamma (NSG) mice, reisolating PDX-cells from mice, PDX cell culture, lentiviral transduction, PDX cell amplification in mice, enrichment of transgenic cells and *in vivo* imaging were described previously (35–37). To save one round of passaging through mice, PDX cells freshly transduced with lentiviruses were kept in culture for 4 days to allow marker expression and enrichment of transgenic cells by flow cytometry before injection into mice.

### *In vivo* assays and Tamoxifen administration

For pairwise competitive *in vivo* experiments, PDX cells transduced with either the control shRNA expressing iRFP (iRFP720) or the shRNA against the GOI expressing mTagBFP were mixed at a 1:1 ratio (shCTRL/shGOI mix) and a total of 300.000 cells were injected into the tail vein of recipient NSG mice. For comparison and to serve as control, a second group of mice was injected, in all the experiments, with the shCTRL/shCTRL mix, consisting of PDX cells transduced with either the control shRNA expressing iRFP or the control shRNA expressing mTagBFP. To promote Cre-ER^*T2*^ translocation to the nucleus and induction of RNA interference, Tamoxifen (TAM, Cat.T5648-5G, Sigma) was resuspended in a sterile mixture of 90% corn oil (Cat.C8267-500ML, Sigma) and 10% ethanol at final concentration of 20mg/ml; aliquots were stored for a maximum of 3 months at -20°C. Before administration to mice, the solution was heated to 37°C and applied orally via gavaging. TAM concentrations were titrated to induce substantial shRNA expression and was given once at 50 mg/kg for AML-388, AML-393, ALL-199, ALL-265 and ALL-811, while animals with samples AML-491 and ALL-707 received 100 mg/kg TAM on two consecutive days.

### Flow cytometric analysis of competitive *in vivo* experiments

Freshly isolated bone marrow cells were analyzed using LSRII (BD Bioscience) to determine fluorochrome distributions. Forward/Side scatter analysis was used to gate on living cells, followed by gating on mCherry (Cre-ER^*T*2^) positive PDX cells. At the beginning, the two cell populations of the mixture were distinguished by expression of either iRFP or mTagBFP. Upon Cre-ER^*T*2^-mediated recombination, cells expressing shCTRL started expressing T-Sapphire (instead of iRFP), while cells expressing shGOI expressed eGFP (instead of mTagBFP) (*Figure S1b*); the color switch was monitored in two separate histograms for either T-Sapphire or eGFP (Figure 1b). The final analysis combined and compared all cells expressing either of the two shRNAs, either T-Sapphire/shCTRL or eGFP/shGOI (Figures 1b and 1c). The distribution of the subpopulations in the shCTRL/shCTRL mixture at 72h after TAM administration was used as reference for analysis, as expression levels of the inducible fluorochromes remained stable after this time point; it enabled controlling for sample-specific, inter-experimental variations in recombination efficiencies and reliably determining the rapid depletion of shGOI expressing cells after TAM, e.g. in *Figure S2a, S2e-f*. To separate shCTRL and shGOI populations for further investigations, cells were sorted using FACSAria III (BD Bioscience).

### Statistical analysis

Statistical significance of pairwise competitive *in vivo* experiments was analyzed by comparing the percentage of eGFP-positive cells out of all recombined cells (sum of T-Sapphire positive plus eGFP positive cells) between the shCTRL/shCTRL mix at 72h after TAM administration with the shCTRL/shGOI mix at the end of each experiment. Statistical analyses were performed using GraphPad Prism 8. Student’s t-test was used, if not differently stated in the legends. A p-value of <0.05 was considered significant.

### *In vivo* drug treatment

For *in vivo* treatment with ABT-199 (Venetoclax, SelleckChem, USA) or Cytarabine, mice were injected with a 1:1 mixture of shCTRL/shMCL1 AML-388 PDX cells (300.000 cells/mouse) and TAM was administered one week thereafter to all animals. 72h after TAM, three mice were sacrificed to determine recombination efficiency. The remaining animals were divided into two groups and treated either with the solvent PBS (n=3) or ABT-199 (100 mg/kg - i.p - 5 consecutive days - 2 cycles - n=3) or Cytarabine (100 mg/kg - i.p - 5 consecutive days - 1 cycle - n=3). At the end of the experiment, mice were sacrificed and BM cells analyzed by flow cytometry for subpopulations’ distribution (see Figure 2h and *Figure S2n*).

### Engraftment of PDX cells in zebrafish

For PDX cell preparation, AML-388 PDX cells expressing (i) mCherry-Cre-ER^*T2*^ and (ii) a knockdown construct, either mTag-BFP/shCTRL or mTagBFP/shMCL1, were amplified in two different donor mice. Mice were sacrificed, human cells isolated and treated *in vitro* with 50nM TAM (Sigma-Aldrich, H7904-25G) to induce recombination and shRNA expression. To allow competitive experiments comparing cells with and without recombination, mCherry positive cells were sorted 48 hours after TAM to gain a 1:1 mixture of eGFP/mtagBFP positive cells and thus 50% of cells with Cre-ER^*T*2^-induced recombination, for either of the two constructs, mTagBFP/shCTRL or mTagBFP/shMCL1. 48 hours post fertilization, dechorionated zebrafish embryos of the Casper lineage (danio rerio) were anesthetized with Tricaine methanesulfonate. Embryos were injected through the Duct of Cuvier, using a Femtojet microinjector (Eppendorf, Hamburg, Germany), with 200 to 500 AML-388 PDX cells per embryo of either of the two mixtures, mTagBFP/shCTRL or mTagBFP/shMCL1. Embryos were raised at 36°C for one day, resulting larvae anesthetized with Tricaine methane-sulfonate and embedded in 1.5% low melting-temperature agarose. For each larva, two fields of view (A and B) were imaged using a spinning disc microscope (20x magnification) and images applied to maximal intensity projection. Using the spot detection function (LoG detector) of the Image-J plugin TrackMate (38) PDX cells were identified by mCherry-Cre-ER^*T2*^ expression and their eGFP intensity determined to quantify the subfraction of cells expressing the shRNA. For each field of view, the ratio between eGFP positive, shRNA expressing cells among all mCherry positive PDX cells was calculated.

### Flow cytometric analysis of BH3 proteins’ level and Annexin V staining

To determine intracellular expression levels of BH3 proteins, cells were fixated in 2% paraformaldehyde, permeabilized using perm/wash buffer (BD Bioscience, Franklin Lakes, NJ, USA) and subsequently stained with fluorescently labeled antibodies against BCL-2 (clone Bcl-2/100, BD Bioscience), BCL-XL (clone 54H11, Cell Signaling, Cambridge, UK), MCL-1 (Clone D2W9E, Cell signaling) or respective isotype controls (Cat.: 556357, BD Bioscience; clone DA1E, Cell Signaling). Dead cells were excluded by Fixable Viability Dye staining. If not otherwise stated, reagents and antibodies were purchased from eBioscience. Flow analysis was performed on a BD FACS Canto II (BD Bioscience) and data were analyzed using FlowJo software (TreeStar Inc., Ashland, OR, USA). Annexin V staining was performed on PDX AML-388 cells isolated from the mouse BM 72h after TAM treatment using PE/Cy7 Annexin V (BioLegend, 640949) according to the manufacturer’s instruction and analyzed by flow cytometry (LSRII, BD Bioscience).

### Targeted genome sequencing

The MLL-AF4 breakpoint was sequenced at the certified laboratory for Leukemia Diagnostics, Department of Medicine III, University Hospital, LMU Munich, Munich, Germany.

### Real-time quantitative PCR

Total RNA from flow cytometry enriched populations was extracted using RNeasy Mini Kit (Qiagen, Venlo, Netherlands) and reverse transcribed using the QuantiTect Reverse Transcription kit (Qiagen, Venlo, Netherlands) according to manufacturer’s instruction. Quantitative PCR was performed in a LightCycler 480 (Roche, Mannheim, Germany) using the corresponding LightCycler 480 Probes Master and the pre-designed Probes of the Universal ProbeLibrary (Roche, Mannheim, Germany). The primer and probes used for qPCR are: HPRT1_fw: tgatagatccattcctatgactgtaga, HPRT1_rv: caagacattctttccagttaaagttg, UPL #22; MLL/AF4_fw: AAGTTCCCAAAACCACTCCTAGT, MLL/AF4 rv: GCCATGAATGGGTCATTTCC, UPL #22; MLL_fw: AAGTTCCCAAAACCACTCCTAGT, MLL_rv: GATCCTGTGGACTCCATCTGC, UPL #22: AF4_fw: CTCCCCTCAAAAAGTGTTGC, AF4_rv: TAGGTCTGCTCAACTGACTGAG, UPL #84. Relative gene expression levels were normalized to HPRT1 using the 2^*−ddCt*^ method.

### Gene expression profiling

Gene expression analysis was performed by applying a bulk adjusted SCRB-Seq protocol on sorted subpopulations from AML PDX samples as described previously (39, 40). All raw fastq data was processed with zUMIs (41) (2.4.5b). Mapping was performed using STAR 2.6.0a (42) against the concatenated human (hg38) and mouse genome (mm10). Gene annotations were obtained from Ensembl (GRCh38.84/GRCm38.75). Analysis of RNA sequencing data followed standard recommendations (43). Statistical analysis was performed using the R 3.6.1 software package (R Core Team, 2019). In case of multiple testing, p-values were adjusted using the Benjamini-Hochberg procedure (FDR-cutoff <0.05). Gene Set Enrichment Analysis (GSEA) using default settings (version 4.0.2) was used for the association of defined gene sets with different subgroups (44).

### Protein immunoassay

To quantify protein of low PDX cell numbers, the Simple Western protein immunoassay (WES, ProteinSimple, San Jose, USA) was performed according to manufacturer’s instructions. Flow cytometry enriched cell populations were incubated in lysis buffer (9803, Cell Signaling Technology, Boston, USA) on ice for 30 min and protein concentration measured by BCA assay (7780, New England Biolabs, Beverly, USA). Results were analyzed using the Compass software (ProteinSimple). Antibodies used were MCL1 (D3CA5, Cell Signaling Technologies), DUX4 (MAB9535, R&D system) and *β*-actin (NB600-501SS, Novus Biologicals). Western blot analysis of PDX ALL-265 was performed as previously described (45).

## ACKNOWLEDGEMENTS

We thank Liliana Mura, Fabian Klein, Maike Fritschle, Annette Frank and Miriam Krekel for excellent technical assistance; Markus Brielmeier and team (Research Unit Comparative Medicine) for animal care services; Binje Vick for establishing PDX AML models; Daniela Senft for correcting the manuscript; Karsten Spiekermann and the LFL laboratory for sequencing the MLL-AF4 breakpoint; Johannes W. Bagnoli, Wolfgang Enard and Helmut Blum for generating SCRB-seq data and Jean Pierre Bourquin and Beat Bornhäuser for providing engrafted sample ALL-265.

## Authors contribution

MC designed and performed experiments, analyzed data and wrote the manuscript, with contributions from KV (MLL-AF4), JV (establishing the technique), MB (DUX4-IGH) and WHL (cloning); JB performed and TH analyzed SCRB-seq data; AA and VB performed zebrafish experiments; VD and PJJ quantified BH3 protein expression; BF and KR designed fluorochrome use; CB, MN and DS provided PDX models; CM, MSS and MB developed cloning strategies; IJ supervised the study, and contributed to experimental design, data analysis and writing the manuscript.

## References

1. S. N. Grobner, B. C. Worst, J. Weischenfeldt, I. Buchhalter, K. Kleinheinz, V. A. Rudneva, P. D. Johann, G. P. Balasubramanian, M. Segura-Wang, S. Brabetz, S. Bender, B. Hutter, D. Sturm, E. Pfaff, D. Hubschmann, G. Zipprich, M. Heinold, J. Eils, C. Lawerenz, S. Erkek, S. Lambo, S. Waszak, C. Blattmann, A. Borkhardt, M. Kuhlen, A. Eggert, S. Fulda, M. Gessler, J. Wegert, R. Kappler, D. Baumhoer, S. Burdach, R. Kirschner-Schwabe, U. Kontny, A. E. Kulozik, D. Lohmann, S. Hettmer, C. Eckert, S. Bielack, M. Nathrath, C. Niemeyer, G. H. Richter, J. Schulte, R. Siebert, F. Westermann, J. J. Molenaar, G. Vassal, H. Witt, Icgc PedBrain-Seq Project, Icgc Mmml-Seq Project, B. Burkhardt, C. P. Kratz, O. Witt, C. M. van Tilburg, C. M. Kramm, G. Fleischhack, U. Dirksen, S. Rutkowski, M. Fruhwald, K. von Hoff, S. Wolf, T. Klingebiel, E. Koscielniak, P. Landgraf, J. Koster, A. C. Resnick, J. Zhang, Y. Liu, X. Zhou, A. J. Waanders, D. A. Zwijnenburg, P. Raman, B. Brors, U. D. Weber, P. A. Northcott, K. W. Pajtler, M. Kool, R. M. Piro, J. O. Korbel, M. Schlesner, R. Eils, D. T. W. Jones, P. Lichter, L. Chavez, M. Zapatka, and S. M. Pfister. ISSNThe landscape of genomic alterations across childhood cancers. Nature, 555(7696):321–327, 2018. ISSN 1476-4687 (Electronic) 0028-0836 (Linking). doi: 10.1038/nature25480.

2. J. G. Moffat, F. Vincent, J. A. Lee, J. Eder, and M. Prunotto. Opportunities and challenges in phenotypic drug discovery: an industry perspective. Nat Rev Drug Discov, 16(8):531–543, 2017. ISSN 1474-1784 (Electronic) 1474-1776 (Linking). doi: 10.1038/nrd.2017.111.

3. M. Hay, D. W. Thomas, J. L. Craighead, C. Economides, and J. Rosenthal. Clinical development success rates for investigational drugs. Nat Biotechnol, 32(1):40–51, 2014. ISSN 1546-1696 (Electronic) 1087-0156 (Linking). doi: 10.1038/nbt.2786.

4. J. W. Scannell and J. Bosley. When quality beats quantity: Decision theory, drug discovery, and the reproducibility crisis. PLoS One, 11(2):e0147215, 2016. ISSN 1932-6203 (Electronic) 1932-6203 (Linking). doi: 10.1371/journal.pone.0147215.

5. F. M. Behan, F. Iorio, G. Picco, E. Goncalves, C. M. Beaver, G. Migliardi, R. Santos, Y. Rao, F. Sassi, M. Pinnelli, R. Ansari, S. Harper, D. A. Jackson, R. McRae, R. Pooley, P. Wilkinson, D. van der Meer, D. Dow, C. Buser-Doepner, A. Bertotti, L. Trusolino, E. A. Stronach, J. Saez-Rodriguez, K. Yusa, and M. J. Garnett. Prioritization of cancer therapeutic targets using crispr-cas9 screens. Nature, 568(7753):511–516, 2019. ISSN 1476-4687 (Electronic) 0028-0836 (Linking). doi: 10.1038/s41586-019-1103-9.

6. E. Zeggini, A. L. Gloyn, A. C. Barton, and L. V. Wain. Translational genomics and precision medicine: Moving from the lab to the clinic. Science, 365(6460):1409–1413, 2019. ISSN 1095-9203 (Electronic) 0036-8075 (Linking). doi: 10.1126/science.aax4588.

7. U. Ben-David, B. Siranosian, G. Ha, H. Tang, Y. Oren, K. Hinohara, C. A. Strathdee, J. Dempster, N. J. Lyons, R. Burns, A. Nag, G. Kugener, B. Cimini, P. Tsvetkov, Y. E. Maruvka, R. O’Rourke, A. Garrity, A. A. Tubelli, P. Bandopadhayay, A. Tsherniak, F. Vazquez, B. Wong, C. Birger, M. Ghandi, A. R. Thorner, J. A. Bittker, M. Meyerson, G. Getz, R. Beroukhim, and T. R. Golub. Genetic and transcriptional evolution alters cancer cell line drug response. Nature, 560(7718):325–330, 2018. ISSN 1476-4687 (Electronic) 0028-0836 (Linking). doi: 10.1038/s41586-018-0409-3.

8. S. J. Engle, L. Blaha, and R. J. Kleiman. Best practices for translational disease modeling using human ipsc-derived neurons. Neuron, 100(4):783–797, 2018. ISSN 1097-4199 (Electronic) 0896-6273 (Linking). doi: 10.1016/j.neuron.2018.10.033.

9. E. Driehuis, A. van Hoeck, K. Moore, S. Kolders, H. E. Francies, M. C. Gulersonmez, E. C. A. Stigter, B. Burgering, V. Geurts, A. Gracanin, G. Bounova, F. H. Morsink, R. Vries, S. Boj, J. van Es, G. J. A. Offerhaus, O. Kranenburg, M. J. Garnett, L. Wessels, E. Cuppen, L. A. A. Brosens, and H. Clevers. Pancreatic cancer organoids recapitulate disease and allow personalized drug screening. Proc Natl Acad Sci U S A, 2019. ISSN 1091-6490 (Electronic) 0027-8424 (Linking). doi: 10.1073/pnas.1911273116.

10. M. Bleijs, M. van de Wetering, H. Clevers, and J. Drost. Xenograft and organoid model systems in cancer research. EMBO J, 38(15):e101654, 2019. ISSN 1460-2075 (Electronic) 0261-4189 (Linking). doi: 10.15252/embj.2019101654.

11. E. C. Townsend, M. A. Murakami, A. Christodoulou, A. L. Christie, J. Koster, T. A. DeSouza, E. A. Morgan, S. P. Kallgren, H. Liu, S. C. Wu, O. Plana, J. Montero, K. E. Stevenson, P. Rao, R. Vadhi, M. Andreeff, P. Armand, K. K. Ballen, P. Barzaghi-Rinaudo, S. Cahill, R. A. Clark, V. G. Cooke, M. S. Davids, D. J. DeAngelo, D. M. Dorfman, H. Eaton, B. L. Ebert, J. Etchin, B. Firestone, D. C. Fisher, A. S. Freedman, I. A. Galinsky, H. Gao, J. S. Garcia, F. Garnache-Ottou, T. A. Graubert, A. Gutierrez, E. Halilovic, M. H. Harris, Z. T. Herbert, S. M. Horwitz, G. Inghirami, A. M. Intlekofer, M. Ito, S. Izraeli, E. D. Jacobsen, C. A. Jacobson, S. Jeay, I. Jeremias, M. A. Kelliher, R. Koch, M. Konopleva, N. Kopp, S. M. Kornblau, A. L. Kung, T. S. Kupper, N. R. LeBoeuf, A. S. LaCasce, E. Lees, L. S. Li, A. T. Look, M. Murakami, M. Muschen, D. Neuberg, S. Y. Ng, O. O. Odejide, S. H. Orkin, R. R. Paquette, A. E. Place, J. E. Roderick, J. A. Ryan, S. E. Sallan, B. Shoji, L. B. Silverman, R. J. Soiffer, D. P. Steensma, K. Stegmaier, R. M. Stone, J. Tamburini, A. R. Thorner, P. van Hummelen, M. Wadleigh, M. Wiesmann, A. P. Weng, J. U. Wuerthner, D. A. Williams, B. M. Wollison, A. A. Lane, A. Letai, M. M. Bertagnolli, J. Ritz, M. Brown, H. Long, J. C. Aster, M. A. Shipp, J. D. Griffin, and D. M. Weinstock. The public repository of xenografts enables discovery and randomized phase ii-like trials in mice. Cancer Cell, 29(4):574–586, 2016. ISSN 1878-3686 (Electronic) 1535-6108 (Linking). doi: 10.1016/j.ccell.2016.03.008.

12. J. G. Clohessy and P. P. Pandolfi. The mouse hospital and its integration in ultra-precision approaches to cancer care. Front Oncol, 8:340, 2018. ISSN 2234-943X (Print) 2234-943X (Linking). doi: 10.3389/fonc.2018.00340.

13. Christopher H. Hulton, Emily A. Costa, Nisargbhai S. Shah, Alvaro Quintanal-Villalonga, Glenn Heller, Elisa de Stanchina, Charles M. Rudin, and John T. Poirier. Direct genome editing of patient-derived xenografts using crispr-cas9 enables rapid in vivo functional genomics. Nature Cancer, 1(3):359–369, 2020. ISSN 2662-1347. doi: 10.1038/s43018-020-0040-8.

14. Y. Koga and A. Ochiai. Systematic review of patient-derived xenograft models for preclinical studies of anti-cancer drugs in solid tumors. Cells, 8(5), 2019. ISSN 2073-4409 (Print) 2073-4409 (Linking). doi: 10.3390/cells8050418.

15. P. Richter-Pechanska, J. B. Kunz, B. Bornhauser, C. von Knebel Doeberitz, T. Rausch, B. Erarslan-Uysal, Y. Assenov, V. Frismantas, B. Marovca, S. M. Waszak, M. Zimmermann, J. Seemann, M. Happich, M. Stanulla, M. Schrappe, G. Cario, G. Escherich, K. Bakharevich, R. Kirschner-Schwabe, C. Eckert, M. U. Muckenthaler, J. O. Korbel, J. P. Bourquin, and A. E. Kulozik. Pdx models recapitulate the genetic and epigenetic landscape of pediatric t-cell leukemia. EMBO Mol Med, 10(12), 2018. ISSN 1757-4684 (Electronic) 1757-4676 (Linking). doi: 10.15252/emmm.201809443.

16. R. Feil, J. Wagner, D. Metzger, and P. Chambon. Regulation of cre recombinase activity by mutated estrogen receptor ligand-binding domains. Biochem Biophys Res Commun, 237 (3):752–7, 1997. ISSN 0006-291X (Print) 0006-291X (Linking). doi: 10.1006/bbrc.1997.7124.

17. F. Schnutgen, N. Doerflinger, C. Calleja, O. Wendling, P. Chambon, and N. B. Ghyselinck. A directional strategy for monitoring cre-mediated recombination at the cellular level in the mouse. Nat Biotechnol, 21(5):562–5, 2003. ISSN 1087-0156 (Print) 1087-0156 (Linking). doi: 10.1038/nbt811.

18. F. Stegmeier, G. Hu, R. J. Rickles, G. J. Hannon, and S. J. Elledge. A lentiviral microrna-based system for single-copy polymerase ii-regulated rna interference in mammalian cells. Proc Natl Acad Sci U S A, 102(37):13212–7, 2005. ISSN 0027-8424 (Print) 0027-8424 (Linking). doi: 10.1073/pnas.0506306102.

19. R. W. Siegel, R. Jain, and A. Bradbury. Using an in vivo phagemid system to identify non-compatible loxp sequences. FEBS Lett, 505(3):467–73, 2001. ISSN 0014-5793 (Print) 0014-5793 (Linking). doi: 10.1016/s0014-5793(01)02806-x.

20. O. Zapata-Hommer and O. Griesbeck. Efficiently folding and circularly permuted variants of the sapphire mutant of gfp. BMC Biotechnol, 3:5, 2003. ISSN 1472-6750 (Electronic) 1472-6750 (Linking). doi: 10.1186/1472-6750-3-5.

21. J. M. Hilden, P. A. Dinndorf, S. O. Meerbaum, H. Sather, D. Villaluna, N. A. Heerema, R. McGlennen, F. O. Smith, W. G. Woods, W. L. Salzer, H. S. Johnstone, Z. Dreyer, G. H. Reaman, and Group Children’s Oncology. Analysis of prognostic factors of acute lymphoblastic leukemia in infants: report on ccg 1953 from the children’s oncology group. Blood, 108(2):441–51, 2006. ISSN 0006-4971 (Print) 0006-4971 (Linking). doi: 10.1182/blood-2005-07-3011.

22. M. Thomas, A. Gessner, H. P. Vornlocher, P. Hadwiger, J. Greil, and O. Heidenreich. Targeting mll-af4 with short interfering rnas inhibits clonogenicity and engraftment of t(4;11)-positive human leukemic cells. Blood, 106(10):3559–66, 2005. ISSN 0006-4971 (Print) 0006-4971 (Linking). doi: 10.1182/blood-2005-03-1283.

23. Y. Fernandez-Marrero, S. Spinner, T. Kaufmann, and P. J. Jost. Survival control of malignant lymphocytes by anti-apoptotic mcl-1. Leukemia, 30(11):2152–2159, 2016. ISSN 1476-5551 (Electronic) 0887-6924 (Linking). doi: 10.1038/leu.2016.213.

24. A. W. Hird and A. E. Tron. Recent advances in the development of mcl-1 inhibitors for cancer therapy. Pharmacol Ther, 198:59–67, 2019. ISSN 1879-016X (Electronic) 0163-7258 (Linking). doi: 10.1016/j.pharmthera.2019.02.007.

25. R. Koch, A. L. Christie, J. L. Crombie, A. C. Palmer, D. Plana, K. Shigemori, S. N. Morrow, A. Van Scoyk, W. Wu, E. A. Brem, J. P. Secrist, L. Drew, A. G. Schuller, J. Cidado, A. Letai, and D. M. Weinstock. Biomarker-driven strategy for mcl1 inhibition in t-cell lymphomas. Blood, 133(6):566–575, 2019. ISSN 1528-0020 (Electronic) 0006-4971 (Linking). doi: 10.1182/blood-2018-07-865527.

26. W. Xiang, C. Y. Yang, and L. Bai. Mcl-1 inhibition in cancer treatment. Onco Targets Ther, 11: 7301–7314, 2018. ISSN 1178-6930 (Print) 1178-6930 (Linking). doi: 10.2147/OTT.S146228.

27. T. Yasuda, S. Tsuzuki, M. Kawazu, F. Hayakawa, S. Kojima, T. Ueno, N. Imoto, S. Kohsaka, A. Kunita, K. Doi, T. Sakura, T. Yujiri, E. Kondo, K. Fujimaki, Y. Ueda, Y. Aoyama, S. Ohtake, J. Takita, E. Sai, M. Taniwaki, M. Kurokawa, S. Morishita, M. Fukayama, H. Kiyoi, Y. Miyazaki, T. Naoe, and H. Mano. Recurrent dux4 fusions in b cell acute lymphoblastic leukemia of adolescents and young adults. Nat Genet, 48(5):569–74, 2016. ISSN 1546-1718 (Electronic) 1061-4036 (Linking). doi: 10.1038/ng.3535.

28. D. Schinnerl, E. Mejstrikova, A. Schumich, M. Zaliova, K. Fortschegger, K. Nebral, A. Attarbaschi, K. Fiser, M. O. Kauer, N. Popitsch, S. Haslinger, A. Inthal, B. Buldini, G. Basso, J. P. Bourquin, G. Gaipa, M. Bruggemann, T. Feuerstein, M. Maurer-Granofszky, R. Panzer-Grumayer, J. Trka, G. Mann, O. A. Haas, O. Hrusak, M. N. Dworzak, and S. Strehl. Cd371 cell surface expression: a unique feature of dux4-rearranged acute lymphoblastic leukemia. Haematologica, 104(8):e352–e355, 2019. ISSN 1592-8721 (Electronic) 0390-6078 (Linking). doi: 10.3324/haematol.2018.214353.

29. Y. Tanaka, M. Kawazu, T. Yasuda, M. Tamura, F. Hayakawa, S. Kojima, T. Ueno, H. Kiyoi, T. Naoe, and H. Mano. Transcriptional activities of dux4 fusions in b-cell acute lymphoblastic leukemia. Haematologica, 103(11):e522–e526, 2018. ISSN 1592-8721 (Electronic) 0390-6078 (Linking). doi: 10.3324/haematol.2017.183152.

30. S. Klaeger, S. Heinzlmeir, M. Wilhelm, H. Polzer, B. Vick, P. A. Koenig, M. Reinecke, B. Ruprecht, S. Petzoldt, C. Meng, J. Zecha, K. Reiter, H. Qiao, D. Helm, H. Koch, M. Schoof, G. Canevari, E. Casale, S. R. Depaolini, A. Feuchtinger, Z. Wu, T. Schmidt, L. Rueckert, W. Becker, J. Huenges, A. K. Garz, B. O. Gohlke, D. P. Zolg, G. Kayser, T. Vooder, R. Preissner, H. Hahne, N. Tonisson, K. Kramer, K. Gotze, F. Bassermann, J. Schlegl, H. C. Ehrlich, S. Aiche, A. Walch, P. A. Greif, S. Schneider, E. R. Felder, J. Ruland, G. Medard, I. Jeremias, K. Spiekermann, and B. Kuster. The target landscape of clinical kinase drugs. Science, 358(6367), 2017. ISSN 1095-9203 (Electronic) 0036-8075 (Linking). doi: 10.1126/science.aan4368.

31. Ryan L. Setten, John J. Rossi, and Si-ping Han. The current state and future directions of rnai-based therapeutics. Nature Reviews Drug Discovery, 18(6):421–446, 2019. ISSN 1474-1784. doi: 10.1038/s41573-019-0017-4.

32. R. Pelossof, L. Fairchild, C. H. Huang, C. Widmer, V. T. Sreedharan, N. Sinha, D. Y. Lai, Y. Guan, P. K. Premsrirut, D. F. Tschaharganeh, T. Hoffmann, V. Thapar, Q. Xiang, R. J. Garippa, G. Ratsch, J. Zuber, S. W. Lowe, C. S. Leslie, and C. Fellmann. Prediction of potent shrnas with a sequential classification algorithm. Nat Biotechnol, 35(4):350–353, 2017. ISSN 1546-1696 (Electronic) 1087-0156 (Linking). doi: 10.1038/nbt.3807.

33. C. Fellmann, T. Hoffmann, V. Sridhar, B. Hopfgartner, M. Muhar, M. Roth, D. Y. Lai, I. A. Barbosa, J. S. Kwon, Y. Guan, N. Sinha, and J. Zuber. An optimized microrna backbone for effective single-copy rnai. Cell Rep, 5(6):1704–13, 2013. ISSN 2211-1247 (Electronic). doi: 10.1016/j.celrep.2013.11.020.

34. Lukas E. Dow, Prem K. Premsrirut, Johannes Zuber, Christof Fellmann, Katherine McJunkin, Cornelius Miething, Youngkyu Park, Ross A. Dickins, Gregory J. Hannon, and Scott W. Lowe. A pipeline for the generation of shrna transgenic mice. Nature Protocols, 7 (2):374–393, 2012. ISSN 1750-2799. doi: 10.1038/nprot.2011.446.

35. S. Ebinger, E. Z. Ozdemir, C. Ziegenhain, S. Tiedt, C. Castro Alves, M. Grunert, M. Dworzak, C. Lutz, V. A. Turati, T. Enver, H. P. Horny, K. Sotlar, S. Parekh, K. Spiekermann, W. Hiddemann, A. Schepers, B. Polzer, S. Kirsch, M. Hoffmann, B. Knapp, J. Hasenauer, H. Pfeifer, R. Panzer-Grumayer, W. Enard, O. Gires, and I. Jeremias. Characterization of rare, dormant, and therapy-resistant cells in acute lymphoblastic leukemia. Cancer Cell, 30(6):849–862, 2016. ISSN 1878-3686 (Electronic) 1535-6108 (Linking). doi: 10.1016/j.ccell.2016.11.002.

36. N. Terziyska, C. Castro Alves, V. Groiss, K. Schneider, K. Farkasova, M. Ogris, E. Wagner, H. Ehrhardt, R. J. Brentjens, U. zur Stadt, M. Horstmann, L. Quintanilla-Martinez, and I. Jeremias. In vivo imaging enables high resolution preclinical trials on patients’ leukemia cells growing in mice. PLoS One, 7(12):e52798, 2012. ISSN 1932-6203 (Electronic) 1932-6203 (Linking). doi: 10.1371/journal.pone.0052798.

37. B. Vick, M. Rothenberg, N. Sandhofer, M. Carlet, C. Finkenzeller, C. Krupka, M. Grunert, A. Trumpp, S. Corbacioglu, M. Ebinger, M. C. Andre, W. Hiddemann, S. Schneider, M. Subklewe, K. H. Metzeler, K. Spiekermann, and I. Jeremias. An advanced preclinical mouse model for acute myeloid leukemia using patients’ cells of various genetic subgroups and in vivo bioluminescence imaging. PLoS One, 10(3):e0120925, 2015. ISSN 1932-6203 (Electronic) 1932-6203 (Linking). doi: 10.1371/journal.pone.0120925.

38. J. Y. Tinevez, N. Perry, J. Schindelin, G. M. Hoopes, G. D. Reynolds, E. Laplantine, S. Y. Bednarek, S. L. Shorte, and K. W. Eliceiri. Trackmate: An open and extensible platform for single-particle tracking. Methods, 115:80–90, 2017. ISSN 1095-9130 (Electronic) 1046-2023 (Linking). doi: 10.1016/j.ymeth.2016.09.016.

39. S. Ebinger, C. Zeller, M. Carlet, D. Senft, J. W. Bagnoli, W. H. Liu, M. Rothenberg-Thurley, W. Enard, K. H. Metzeler, T. Herold, K. Spiekermann, B. Vick, and I. Jeremias. Plasticity in growth behavior of patients’ acute myeloid leukemia stem cells growing in mice. Haematologica, 2020. ISSN 1592-8721 (Electronic) 0390-6078 (Linking). doi: 10.3324/haematol.2019.226282.

40. Magali Soumillon, Davide Cacchiarelli, Stefan Semrau, Alexander van Oudenaarden, and Tarjei S. Mikkelsen. Characterization of directed differentiation by high-throughput single-cell rna-seq. bioRxiv, page 003236, 2014. doi: 10.1101/003236.

41. S. Parekh, C. Ziegenhain, B. Vieth, W. Enard, and I. Hellmann. zumis - a fast and flexible pipeline to process rna sequencing data with umis. Gigascience, 7(6), 2018. ISSN 2047-217X (Electronic) 2047-217X (Linking). doi: 10.1093/gigascience/giy059.

42. A. Dobin, C. A. Davis, F. Schlesinger, J. Drenkow, C. Zaleski, S. Jha, P. Batut, M. Chaisson, and T. R. Gingeras. Star: ultrafast universal rna-seq aligner. Bioinformatics, 29(1):15–21, 2013. ISSN 1367-4811 (Electronic) 1367-4803 (Linking). doi: 10.1093/bioinformatics/bts635.

43. T. Herold, V. Jurinovic, A. M. N. Batcha, S. A. Bamopoulos, M. Rothenberg-Thurley, B. Ksienzyk, L. Hartmann, P. A. Greif, J. Phillippou-Massier, S. Krebs, H. Blum, S. Amler, S. Schneider, N. Konstandin, M. C. Sauerland, D. Gorlich, W. E. Berdel, B. J. Wormann, J. Tischer, M. Subklewe, S. K. Bohlander, J. Braess, W. Hiddemann, K. H. Metzeler, U. Mansmann, and K. Spiekermann. A 29-gene and cytogenetic score for the prediction of resistance to induction treatment in acute myeloid leukemia. Haematologica, 103(3):456–465, 2018. ISSN 1592-8721 (Electronic) 0390-6078 (Linking). doi: 10.3324/haematol.2017.178442.

44. V. K. Mootha, C. M. Lindgren, K. F. Eriksson, A. Subramanian, S. Sihag, J. Lehar, P. Puigserver, E. Carlsson, M. Ridderstrale, E. Laurila, N. Houstis, M. J. Daly, N. Patterson, J. P. Mesirov, T. R. Golub, P. Tamayo, B. Spiegelman, E. S. Lander, J. N. Hirschhorn, D. Altshuler, and L. C. Groop. Pgc-1alpha-responsive genes involved in oxidative phosphorylation are coordinately downregulated in human diabetes. Nat Genet, 34(3):267–73, 2003. ISSN 1061-4036 (Print) 1061-4036 (Linking). doi: 10.1038/ng1180.

45. H. Ehrhardt, S. Pfeiffer, D. Schrembs, F. Wachter, M. Grunert, and I. Jeremias. Activation of dna damage response by antitumor therapy counteracts the activity of vinca alkaloids. Anticancer Res, 33(12):5273–87, 2013. ISSN 1791-7530 (Electronic) 0250-7005 (Linking).

