## Supplemental Information for "*In vivo* inducible reverse genetics in patients’ tumors to identify individual therapeutic targets"

<u>This file contains</u>:

Supplemental Tables

Supplemental Figures and Legends

### Table S1. Clinical characteristics of AML and ALL patients

| sample | disease stage* | age [years] | sex | cytogenetics | mutations∞ | mean passaging time§ [days] | Reference |
|---|---|---|---|---|---|---|---|
| **AML-388** | ID | 57 | m | MLL-AF6 | KRAS, CEBPZ | 47 | 1, 11 |
| **AML-393** | R1 | 47 | f | MLL-AF10 | BCOR, KRAS | 49 | 1, 2, 5, 10, 11 |
| **AML-491** | R1 | 53 | f | del(7) (q2?1) | DNMT3A, BCOR, NRAS, KRAS, ETV6, PTPN11, RUNX1 | 59 | 1, 4, 5, 6, 8, 9, 10, 11 |
| **ALL-199** | R2 | 8 | f | somatic trisomy21; leukemic homozygous 9p deletion; P2RY8-CRLF2 | N.D. | 42 | 3, 7 |
| **ALL-265** | R1 | 5 | f | hyperdipoidy with additional 6,13,14,17,18,21,X chromosome | KMT2D, HERC1 | 43 | 3, 7 |
| **ALL-707** | ID | 2 | m | MLL-AF4 | N.D. | 50 | 7 |
| **ALL-811** | R1 | 69 | | DUX4-IGH | N.D. | 80 | |

*when the primary sample was obtained; ∞ mutations determined by panel sequencing; §time of passaging through mice refers to the time from injection of the sample until mice had to be sacrificed due to end stage leukemia; ID = initial diagnosis; R1 = 1st relapse; R2 = 2nd relapse; f = female; m = male; N.D. not determined.


[1] Vick et al., PLoS One 2015
[2] Sandhöfer et al., Leukemia 2015
[3] Ebinger et al., Cancer Cell 2016
[4] Reiter et al., Leukemia 2017
[5] Tzelepis et al., Nature Communications 2018
[6] Garg et al., Blood 2019
[7] Heckl et al., Leuk Lymphoma 2019
[8] Lynch et al., Leukemia 2019
[9] Ruzicka Leukemia 2019
[10] Stief et al., Leukemia 2020
[11] Ebinger et al., Hematologica 2020

### Table S2. shRNA sequences

| target | guide<br>22mer |
|---|---|
| **MLL-AF4** | TGGAGTAGGTCTGCTTTTCTTT |
| | TAGGTCTGCTTTTCTTTTGGTT * |
| **MCL1** | TTACACATCAATTCGTTCTGTA |
| | TGAAACTGAACTTTGCTTCTTT * |
| **DUX4-IGH** | TTCTGAAACCAAATCTGGACCC |
| | TTCGATTCTGAAACCAGATCTG * |

* For each target an additional shRNA sequence was tested.

### Appendix to Table S2.

Sequence of the 110bps oligo to be cloned into the pCDH-plasmid digested with XhoI and EcoRI enzymes:

XhoI — passanger strand — guide strand — EcoRI

TCGAG**AAGGTATATTGCTGTTGACAGTGAGCG**CAAGAAAAGCAGACCTACTCCA**TAGTGAAGCCACAGATGTA**TGGAGTAGGTCTGCTTTTCTTT**TGCCTACTGCCTCG**G

5‘ common flank — loop — MLL/AF4 shRNA sequence — 3‘ common flank

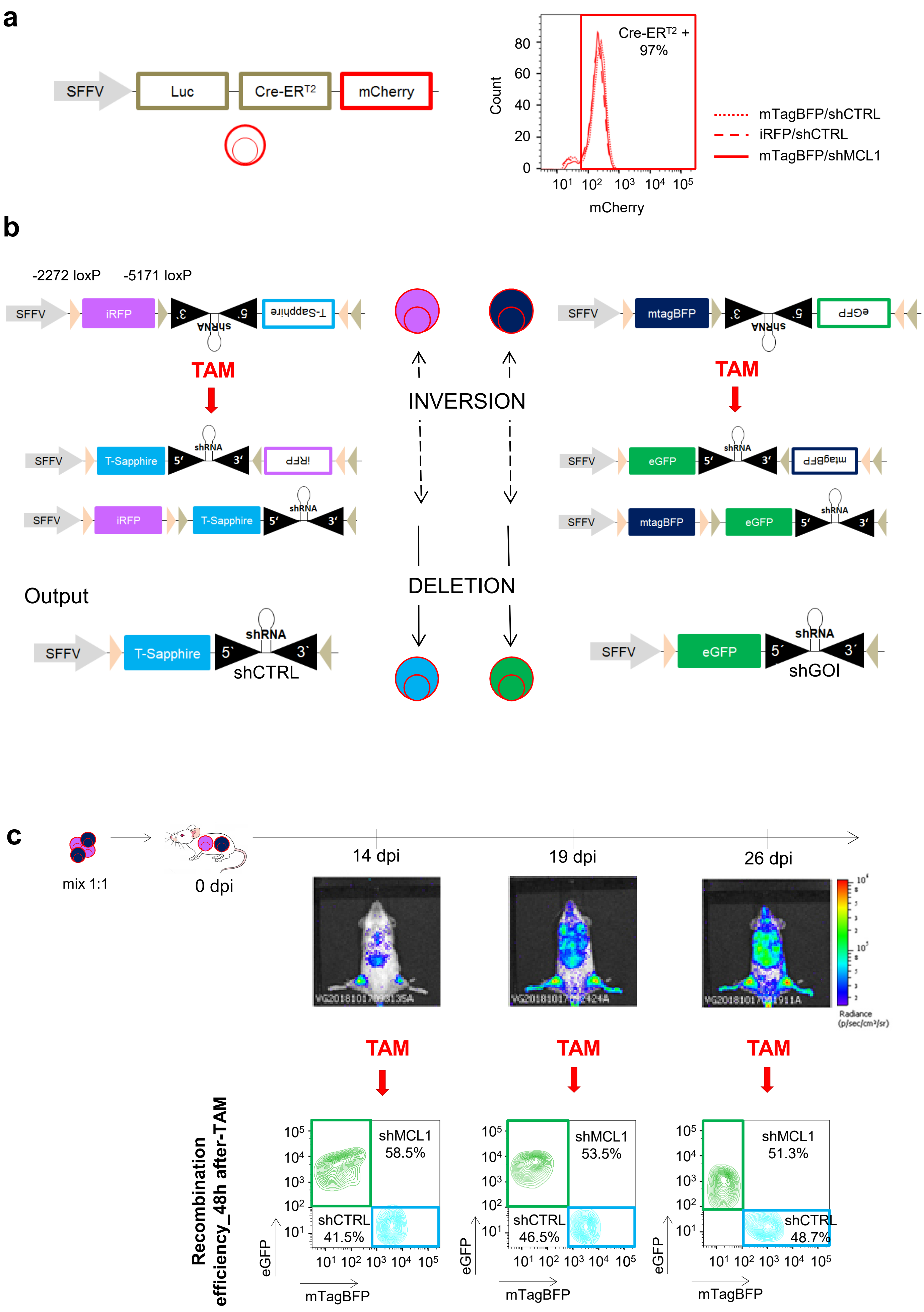


**Suppl. Figure 1**

d

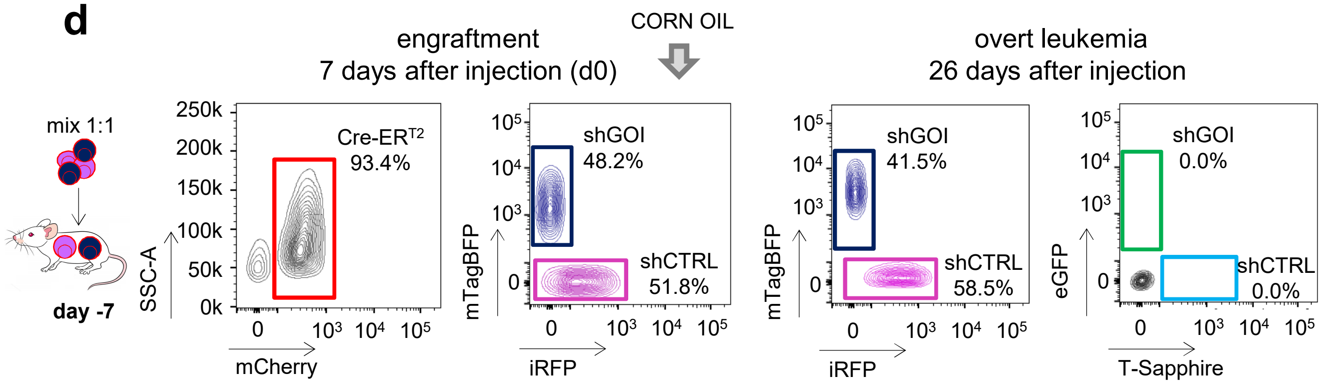


e

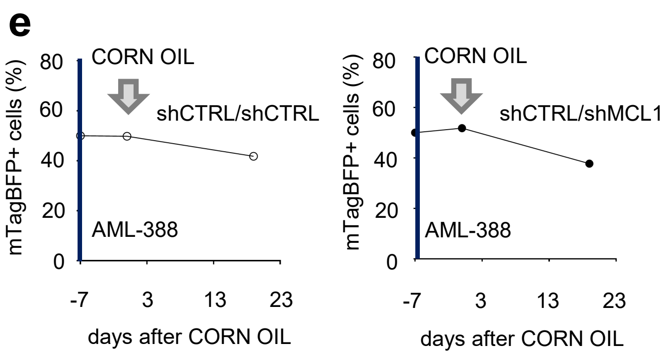


f

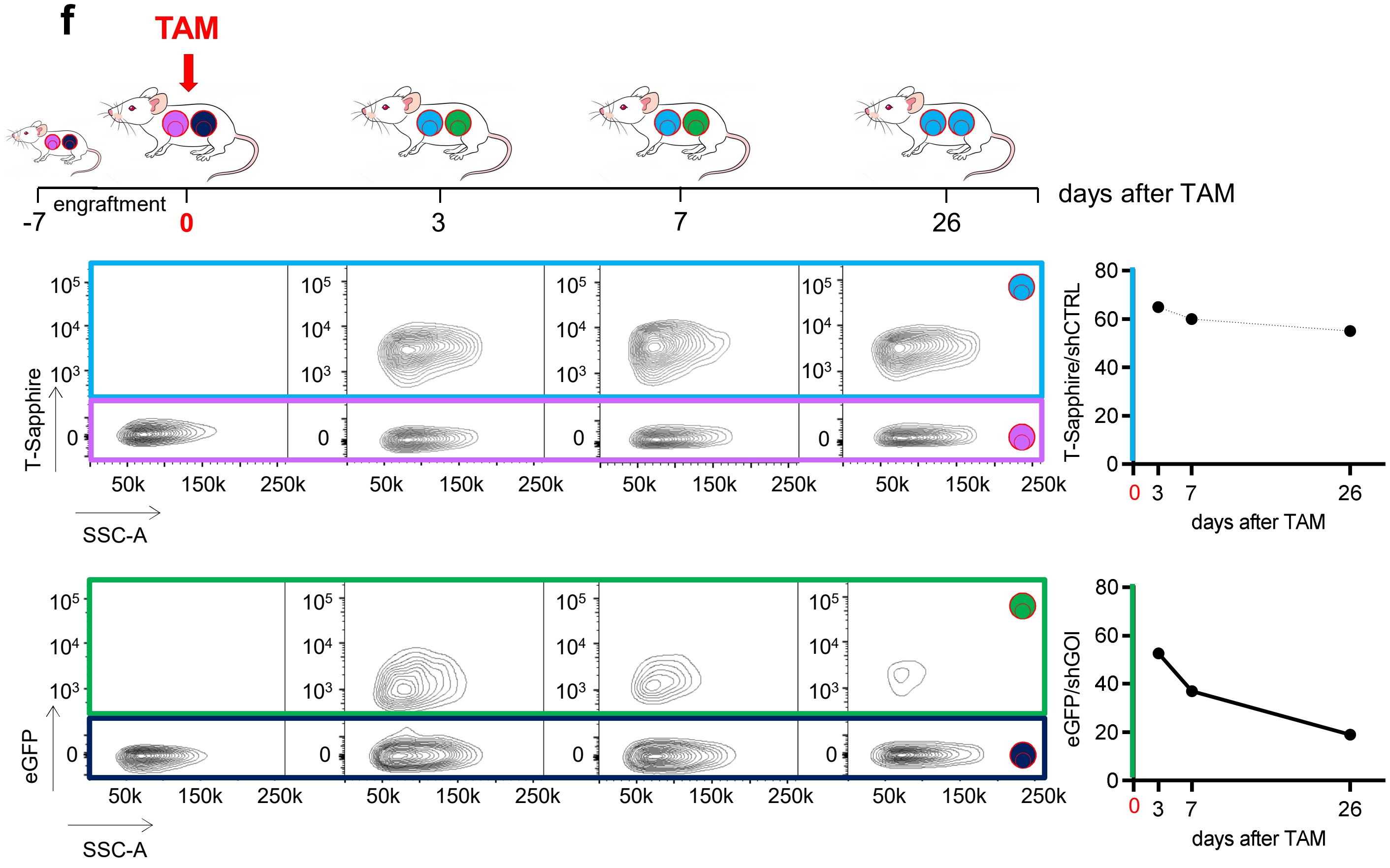


Suppl. Figure 1

**Supplementary Figure 1**

**Inducible knockdown system in PDX acute leukemia models *in vivo* and quality controls**

a) Details of the Cre-ER$^{T2}$ expression construct (left). Expression of a Gaussia luciferase (Luc) for *in vivo* imaging, Cre-ER$^{T2}$ and mCherry are under the control the SFFV promoter and connected via 2A-peptides. Histogram (right) displays expression levels of mCherry in different AML-388 PDX derivatives, co-transduced with different knockdown constructs; similar data were obtained in all 7 PDX models studied.

b) The 2-step process of Cre-ER$^{T2}$ mediated recombination. The shRNA cassette is flanked by two different pairs of loxP sites; upon treatment of mice with TAM, Cre-ER$^{T2}$ translocates to the nucleus and first induces a reversible inversion between either of the two pairs of loxP sites (one example is shown); this converts the out-of-frame cassette into frame so that both, the inducible fluorochrome (T-Sapphire or eGFP) and the coupled shRNA, get under control of the SFFV promoter. In a second step, Cre-ER$^{T2}$ mediates an irreversible deletion between the second pair of loxP sites, resulting in deletion of the original fluorochrome (iRFP or mTagBFP). As end product, the constitutively expressed fluorochrome is lost, while the inducible fluorochrome is expressed in equimolar amounts together with the shRNA.

c) Recombination efficiency is independent from tumor burden. Mice were injected with a mix of shCTRL/shMCL1 cells. Tumor growth was monitored by *in vivo* imaging; at the indicated time points, TAM was administered at 50 mg/kg per mouse to induce Cre-ER$^{T2}$ -mediated inversion/deletion and consequent shRNA expression. Recombination efficiency was analyzed 48h after TAM by quantifying expression of the inducible fluorochrome markers by flow cytometry. Data from representative mice are displayed; 2 mice per time point were analyzed.

d-e) Quality controls in the absence of TAM:

d) Competitive experiments were set up as described in Figure 1c, except that mice were injected with the solvent corn oil alone without TAM. One week after injection (day 0) two mice were sacrificed; flow cytometry shows results from one representative mouse per time point; percentage of iRFP/shCTRL positive versus mTagBFP/shGOI (shMCL1) positive cells was determined from all mCherry-Cre-ER$^{T2}$-positive cells. Corn oil was administered to the remaining two mice and cells analyzed 26 days after by flow cytometry.

e) Percentage of mTagBFP positive cells was quantified from all isolated cells, expressing either mTagBFP or iRFP, for the two different mixtures shCTRL/shCTRL or shCTRL/shMCL1.

f) Data complementing Figure 1c; from the shCTRL/shMCL1 mixture, shCTRL cells and shMCL1 cells were analyzed separately and not as pairwise competitive analysis as in Figure 1c. Upper row shows cells harboring the iRFP/shCTRL construct without shCTRL expression converting upon TAM treatment into T-Sapphire/shCTRL with shCTRL expression; lower row shows cells harboring the mTagBFP/shMCL1 construct without shMCL1 expression converting into eGFP/shMCL1 with shMCL1 expression. Right panels show quantification as [eGFP/shGOI positive cells divided by (the sum of mTagBFP/shGOI positive plus eGFP/shGOI positive cells)], respectively. The reliability of this type of analysis is restricted to settings with low cell death within the first 3 days.

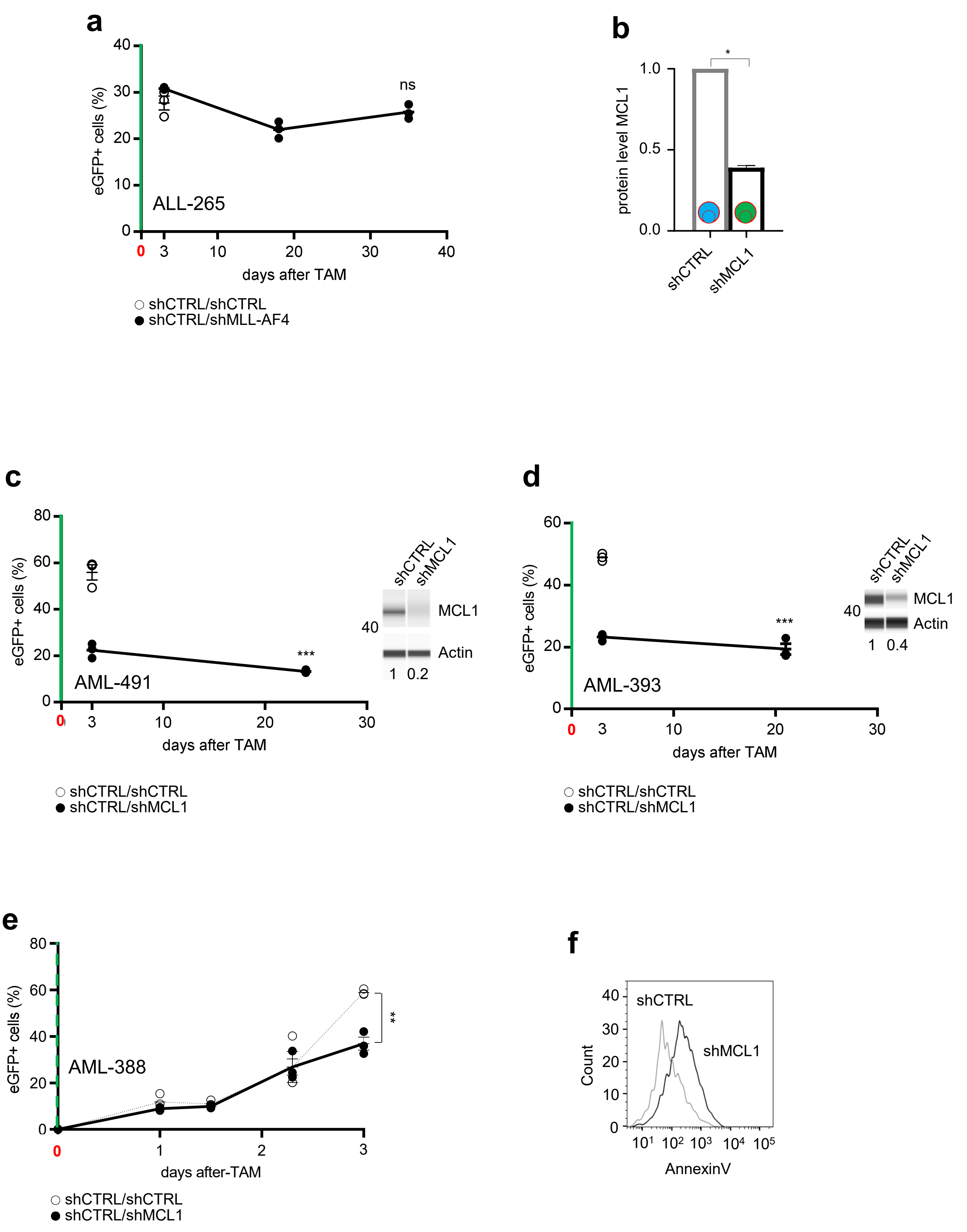


**Suppl. Figure 2**

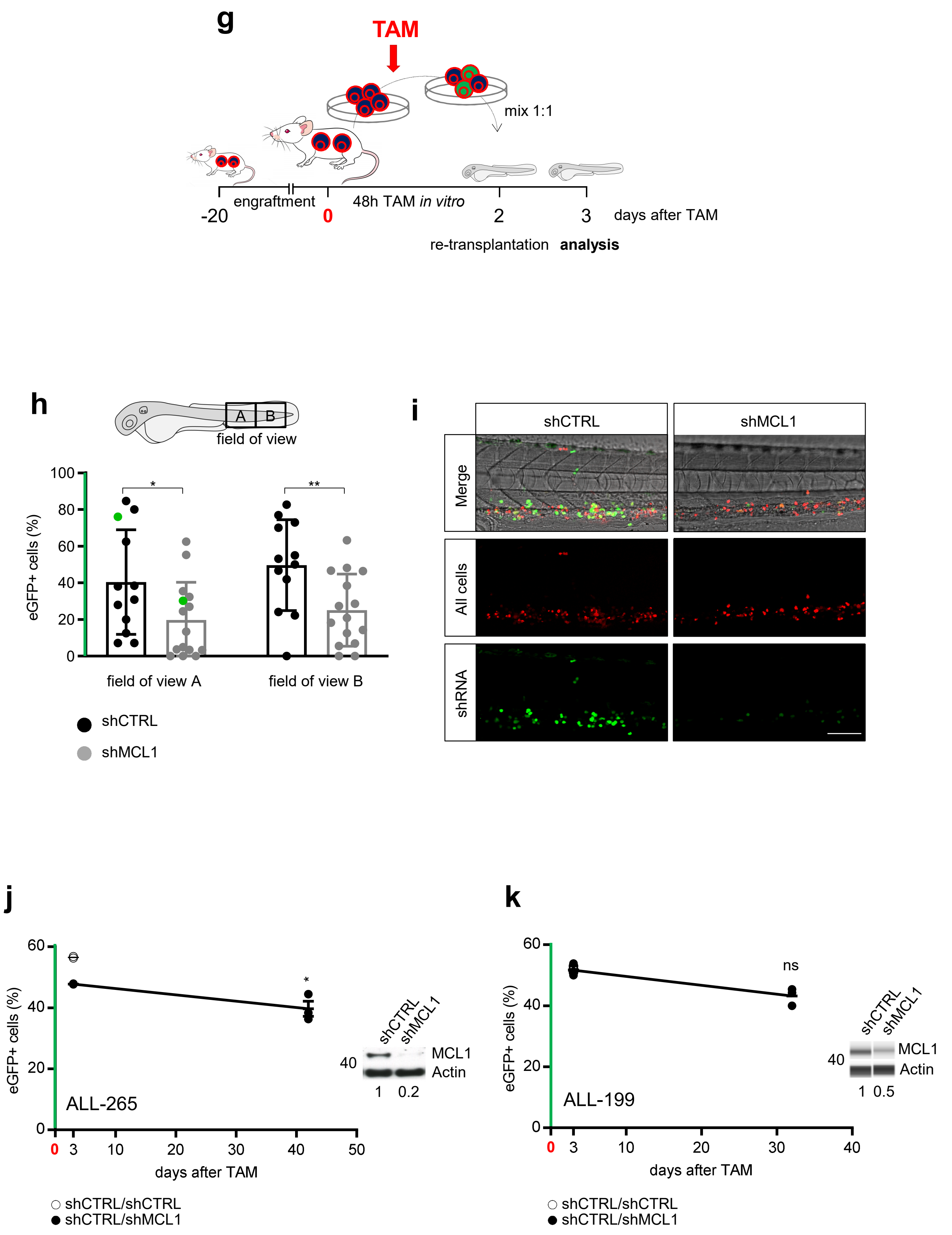


**Suppl. Figure 2**

**l**

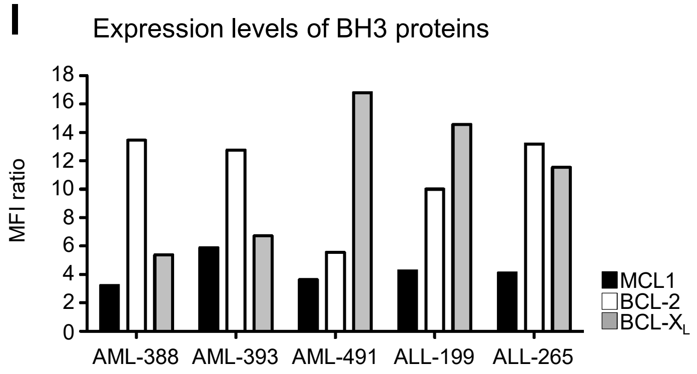
Expression levels of BH3 proteins
MFI ratio
18
16
14
12
10
8
6
4
2
0
AML-388
AML-393
AML-491
ALL-199
ALL-265
MCL1
BCL-2
BCL-XL

**m**

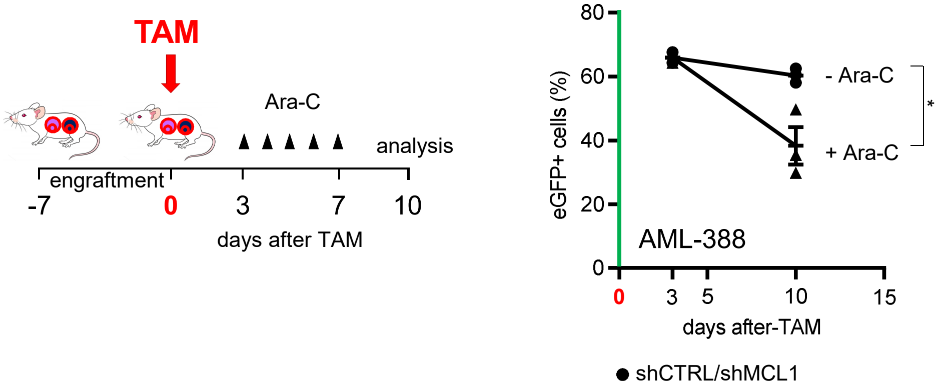
TAM
Ara-C
analysis
engraftment
-7
0
3
7
10
days after TAM
Treatment with Cytarabine
eGFP+ cells (%)
80
60
40
20
0
- Ara-C
+ Ara-C
*
AML-388
0
3
5
10
15
days after-TAM
shCTRL/shMCL1

**n**

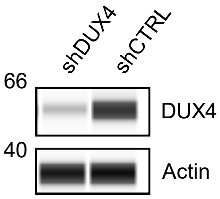
shDUX4
shCTRL
66
40
DUX4
Actin

**Supplementary Figure 2** *In vivo* **functional validation of essential genes**

a) Experiment described in Figure 2b was performed using the non MLL-AF4 rearranged ALL-265 PDX as control. Each dot represents one mouse (n=3 for shCTRL/shCTRL; n=6 for the shCTRL/shMCL1), mean ± SEM, unpaired t-test.

b) Quantification of MCL1 protein expression as mean from 3 independent experiments, as described in Figure 2e; mean ± SD is shown; * p≤0.05 by t-test.

c-d) Raw data of experiments quantified in Figure 2g. Experiment described in Figure 2d was identically performed and depicted in 2 additional AML samples. Mean ± SEM of results for AML-491 (c), AML-393 (d) is shown. At the end of each experiment, MCL1 protein expression was analyzed in sorted shCTRL and shMCL1 populations by protein immunoassay. ***p<0.001, **p<0.01, ns not significant by unpaired t-test.

e) MCL1 knockdown cells are depleted early after TAM induction. For a kinetic of eGFP-expression at early time points after TAM, competitive experiments were performed and each subpopulation analyzed separately as described in Figure S1f; mice were analyzed at 24, 36, 52 and 72 hours after TAM administration. The analysis shows quantification as [eGFP/shMCL1 positive cells divided by (the sum of mTagBFP/shMCL1-positive plus eGFP/shMCL1-positive cells)]. The same analysis was performed for the shCTRL/shCTRL mixture. Mean ± SEM of 3 mice per group per time point is displayed. ** ≤0,01, by unpaired t-test.

f) Knockdown of MCL1 induces apoptosis; Annexin V staining in PDX AML-388 72h after TAM. Representative histograms of 3 experiments are shown.

g-i) Transplantation of MCL1 knockdown PDX cells in zebrafish.

g) Experimental layout: mCherry-Cre-ER$^{T2}$ positive PDX cells from donor mice injected with AML-388 mTagBFP/shCTRL or mTagBFP/shMCL1 cells were isolated from the BM of mice 20 days after injection. Cells were treated *in vitro* with 50 nM TAM to induce eGFP/shRNA expression. After 48 hours, PDX cells were sorted to adjust cells with (eGFP positive) and without recombination (mTagBFP positive) to a 1:1 ratio. Cells were re-transplanted into groups of zebrafish embryos at 48 hours post fertilization at 200 to 500 PDX cells per embryo. 24 hours post transplantation (72h after TAM), larvae were anesthetized and two fields of view (A and B) of the caudal hematopoietic tissue of each larvae were imaged to quantify mCherry and eGFP-positive cells.

h) Mean ± SEM of the percentage of eGFP/shCTRL (n=12) and eGFP/shMCL1 (n=15) positive cells among all transplanted, mCherry positive cells is shown. In both fields, each dot represents one fish, green dots represent the value of the images displayed in (i). * p<0.05, ** p<0.01, by unpaired t-test.

i) Images of representative larvae injected with eGFP/shCTRL (left) or eGFP/shMCL1 (right) expressing cells are displayed. Upper panel depicts merged images of the brightfield shot for anatomic orientation; mCherry positive cells are shown in red in the middle panel and eGFP/shRNA positive cells are shown in green in lower panel. Scale bar 100µm.

j-k) Raw data of experiments quantified in Figure 2g. Experiment described in Figure 2d was identically performed and depicted in 2 additional ALL samples. Mean ± SEM of results for ALL-265 (j) and ALL-199 (k) is shown. At the end of each experiment, MCL1 protein expression was analyzed in sorted shCTRL and shMCL1 populations by protein immunoassay (ALL-199) or traditional Western blot (ALL-265). ***p<0.001, **p<0.01, ns not significant by unpaired t-test

l) Intracellular expression levels of MCL1, BLC-2 and BCL-$X_L$, as measured by flow cytometry in the indicated PDX samples. Protein expression was calculated as the ratio of stained antibody mean fluorescent intensity (MFI) divided by isotype control MFI.

m) shCTRL/shMCL1 expressing AML-388 cells were studied and the experiment set up and analyzed as in Figure 2h, except that Cytarabine was administered 3 days after TAM treatment at a dose of 100 mg/kg/per day i.p. for 5 consecutive days. At the end of the experiment (10 days post-TAM), mice were analyzed as in Figure 1c. Mean ± SEM of 3 replicates per group and condition are shown. *p<0.05 by unpaired t-test.

n) Protein immunoassay of DUX4 in NALM-6 cells, after lentiviral transduction with the indicated shRNAs. β-actin was used as loading control.
